# TRIC Coupled with TR-FRET as a High-Throughput Screening Platform for the Discovery of SLIT2 Binders: A Proof-of-Concept Approach

**DOI:** 10.1101/2025.07.08.663693

**Authors:** Nelson García-Vázquez, Moustafa T. Gabr

**Affiliations:** Department of Radiology, Molecular Imaging Innovations Institute (MI3), Weill Cornell Medicine, New York, NY 10065, USA

**Keywords:** SLIT2, TRIC, Small Molecules, High-Throughput Screening, Drug Discovery, Assay Development

## Abstract

SLIT2, a secreted glycoprotein involved in axon guidance, immune modulation, and tumor progression, remains largely unexplored as a pharmacological target due to the absence of small-molecule modulators. Here, we present a proof-of-concept high-throughput screening platform that integrates Temperature-Related Intensity Change (TRIC) technology with time-resolved Förster resonance energy transfer (TR-FRET) to identify small molecules capable of disrupting the SLIT2/ROBO1 interaction. Screening a lipid metabolism–focused compound library (653 molecules) yielded bexarotene, as the most potent small molecule SLIT2 binder reported to date, with a dissociation constant (*K*_D_) of 2.62 µM. Follow-up TR-FRET assays demonstrated dose-dependent inhibition of SLIT2/ROBO1 interaction, with an IC_50_ value of ∼22.8 µM and maximal inhibition of ∼15– 25%. These findings suggest a novel extracellular activity of bexarotene and validate the combined use of TRIC and TR-FRET as a scalable screening strategy for SLIT2-targeted small molecules. This platform lays the groundwork for future high-throughput discovery efforts against SLIT2 and its signaling axis.

**Graphical abstract:** TRIC-based small molecule screening platform protocol steps with implementation of TR-FRET for the identification of SLIT2 inhibitors.

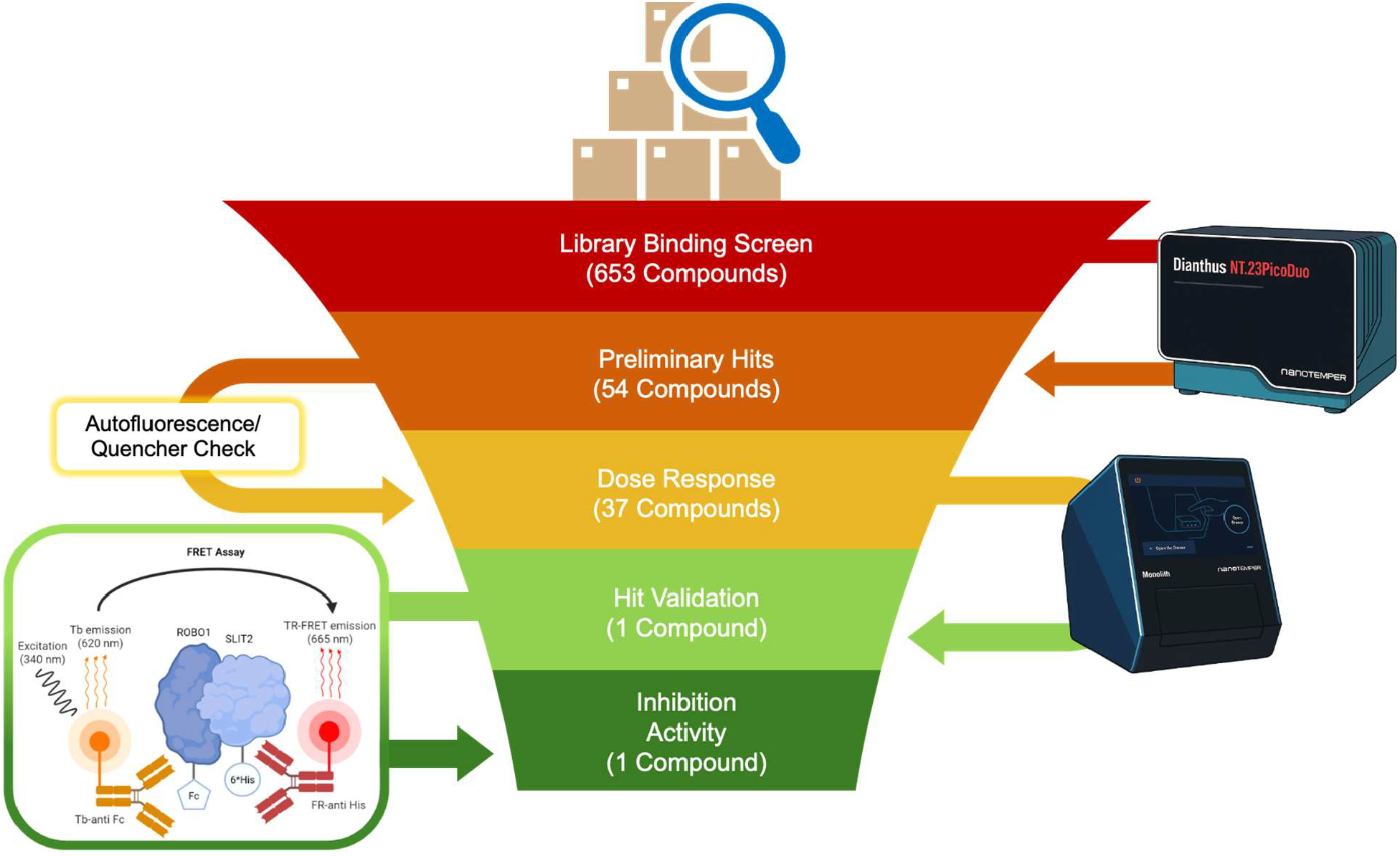

## 1. Introduction

SLIT2 is a large, secreted glycoprotein initially identified for its role in repulsive axon guidance during neural development, mediated through interactions with ROBO family receptors. This interaction initiates a cascade of intracellular events that affect cytoskeletal dynamics, cell adhesion, and migration, largely through modulation of Rho GTPase signaling (1,2). Beyond its neural functions, SLIT–ROBO signaling plays pivotal roles in diverse physiological and pathological contexts, such as vascular development, organ morphogenesis, immune cell migration, and tumor progression (3-5).

SLIT2 has been shown to inhibit leukocyte chemotaxis by interfering with chemokine-driven actin remodeling and polarization, thereby suppressing the directional migration of neutrophils, dendritic cells, and T cells (6,7). This inhibitory effect is thought to be mediated through disruption of integrin signaling and actin cytoskeletal dynamics, ultimately limiting leukocyte infiltration into inflamed or damaged tissues. Physiologically, SLIT2 expression is upregulated during tissue repair, where it may contribute to the resolution of inflammation by restraining excessive immune cell recruitment (8). However, this anti-inflammatory function can be co-opted in pathological settings. In tumors, for instance, SLIT2 expression by stromal or tumor-associated cells has been associated with immune evasion. In pancreatic ductal adenocarcinoma and certain gliomas, elevated SLIT2 levels correlate with immunosuppressive microenvironments, reduced infiltration of cytotoxic T cells, and skewing of macrophages toward an M2-like, tumor-promoting phenotype (9,10). These effects appear to be primarily mediated through ROBO1, although ROBO2 may also contribute in a context-dependent manner.

SLIT2 also exhibits tumor-suppressive activity in various epithelial malignancies, where it is frequently downregulated through promoter hypermethylation (11–13). Restoration of SLIT2 expression in these models has been shown to suppress tumor cell migration and invasion, inhibit angiogenesis, and enhance sensitivity to chemotherapy (12,14). While this dual role may appear contradictory, such functional versatility is not uncommon among extracellular signaling proteins. These context-dependent effects likely reflect differences in receptor expression, proteolytic processing, tissue microenvironment, and the specific SLIT2 isoforms or fragments involved—particularly as the full-length protein and its N-terminal fragment may exert distinct, and in some cases opposing, biological activities (15).

Given its context-dependent roles in cell migration, immune regulation, and tissue architecture, SLIT2 represents a compelling yet underexplored therapeutic target. However, the absence of small-molecule probes capable of modulating SLIT2/ROBO signaling has hindered its pharmacological interrogation. Current strategies primarily rely on genetic manipulation or antibody-based inhibition of ROBO1, which lack tunability and temporal precision (16). A small molecule screening approach targeting SLIT2 not only holds promise for identifying novel modulators of this pathway but also provides a platform for the development of scalable and robust discovery assays.

Here, we present a scalable screening platform that integrates Temperature-Related Intensity Change (TRIC) technology with time-resolved Förster resonance energy transfer (TR-FRET) to identify small-molecule binders of SLIT2. Using this dual approach, we identified bexarotene as a small molecule SLIT2 binder that inhibits SLIT2/ROBO1 interaction.

## 2. Methods

### Library screening with Dianthus

The small molecule library screen was achieved using the TRIC technology on a Dianthus (NanoTemper, Munich, Germany). The Lipid Metabolism Compound Library (653 compounds, #L2510) was purchased from TargetMol (Boston, MA, USA). hSLIT2-His was purchased from AcroByosistems (Cat. No. SL2-H52H6) and was labeled with RED-tris-NTA 2^nd^ generation dye (NanoTemper, #MO-L018) following the manufacturer’s instructions.

Compounds from the library were transferred from the stock 384-well plates to intermediate plates at 2-times the final concentration (200 µM) in a 10mM HEPES, pH 7.4, 0.005% Tween20, 4% DMSO buffer using an Integra mini-96 pipetting tool **(Figure 1, step A)**. 10 µL from this intermediate plate was transferred to Dianthus 384-well plates and incubated for 20 minutes with 50 nM of hSLIT2-His, labeled as described above, resulting in a final compound concentration 100 µM, and final 2% DMSO. The first and last two columns were reference wells composed of labeled hSLIT2-His protein incubated with DMSO. The normalized fluorescence (F_norm_) for all wells were analyzed with GraphPad Prism 10 and those outside of 3 standard deviations from the average of the reference were catalogued as potential hits **(Figure 1, step B)**. Molecules flagged by the instrument’s software (DI.Control) as scan anomalies or aggregation error were excluded.

**Figure 1.**
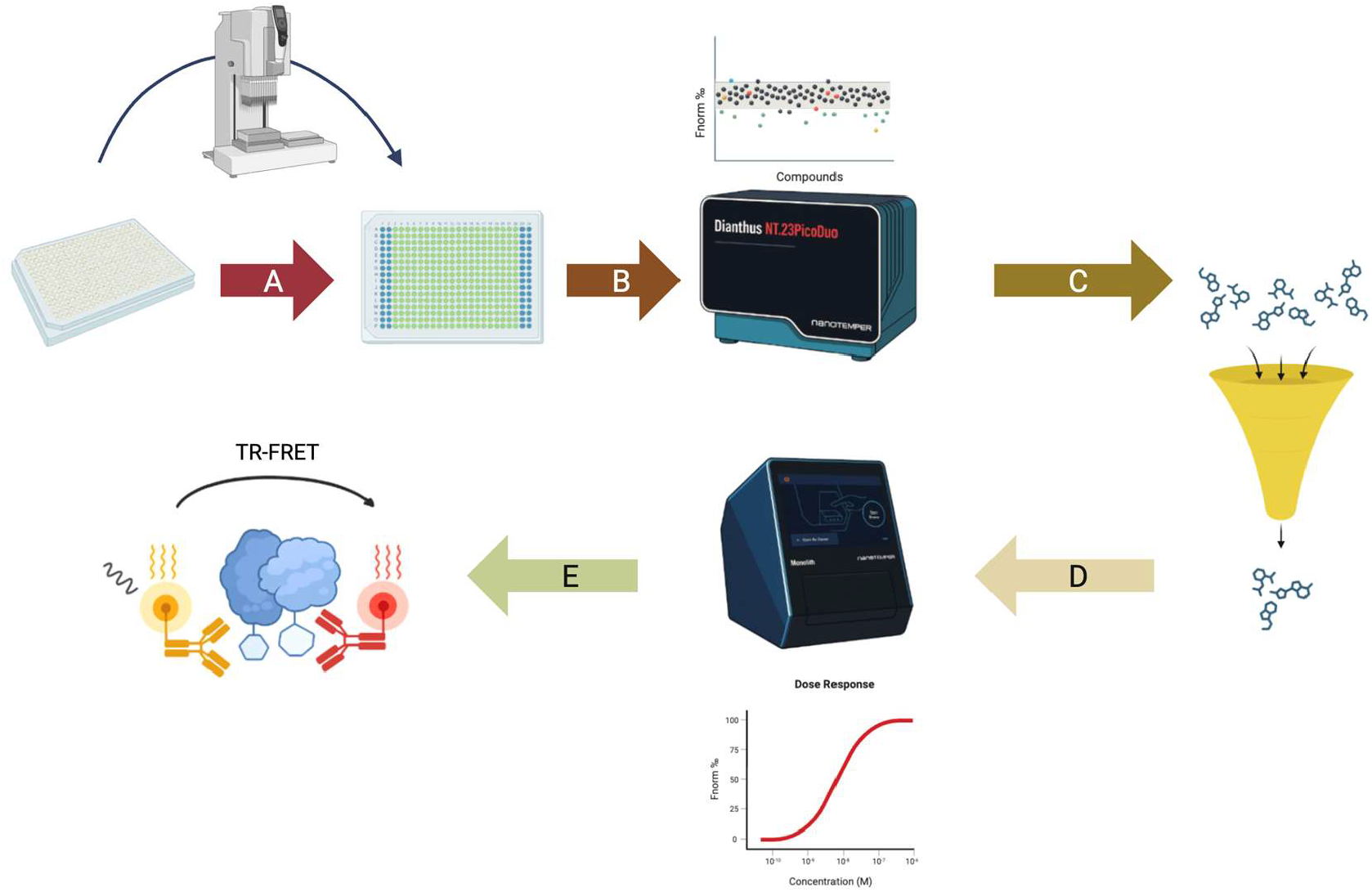
Schematic of assay pipeline. **(A)** Intermediate plate preparation from stock solutions, into the final plate. **(B)** The library is screened using the Dianthus TRIC technology. **(C)** Resulting hits are filtered for potential auto-fluorescence or signal quenching. **(D)** Filtered hits are screened for dose response using a Monolith X. **(E)** Compounds that showcase dose response are screened with a TR-FRET to evaluate SLIT2/ROBO1 inhibition profiles.

### Auto fluorescence/Quencher Filtering

As certain fluorescent compounds may influence the F_norm_ values, the resulting potential hits were filtered to exclude auto fluorescent compounds and signal quenchers **(Figure 1, step C)**. The initial fluorescence for all wells were analyzed with GraphPad Prism 10. Compounds found to have an initial fluorescence signal higher than one standard deviation from the reference average were labeled auto fluorescent, those with lower than one standard deviation were labeled quenchers.

### Binding affinity with Monolith X

Binding affinity tests were accomplished on a Monolith X (NanoTemper, Munich, Germany). The ligands were prepared in 24 assay points in 2-fold dilution series starting from 400 µM (12 points) and 300 µM (12 points), followed by incubation with 50 nM hSLIT2-His, labeled with RED-tris-NTA 2^nd^ generation, for 20 minutes. TRIC/MST measurements were performed at 100% laser excitation and medium power. *K*_D_ values and graphs were calculated and prepared with Graphpad Prism 10 using the F_norm_ values resulting from three independent measurements **(Figure 1, step D)**.

### SLIT2 inhibition with TR-FRET

TR-FRET assay was performed as we previously reported (19). **(Figure 1, step E)**,

## 3. Results

As a preliminary screen, we selected the Lipid Metabolism Library from TargetMol. Although SLIT2 is classically associated with axon guidance and cell migration, its signaling through ROBO receptors—particularly via Rho GTPases such as Rac1 and Cdc42 (1,2)—is tightly linked to membrane organization and lipid-mediated signaling pathways. These interactions occur at the cell surface and are modulated by membrane composition (3). Moreover, SLIT2 has been shown to suppress NF-κB activity in neutrophils and inhibit inflammatory cell migration (6), actions reminiscent of lipid-derived mediators. While SLIT2 is not directly involved in lipid metabolism, its signaling intersects with lipid-regulated processes, supporting the rationale for screening lipid-targeting compounds as potential functional modulators.

For the initial single-dose screening, compounds were classified as normal, fluorescence quenchers, or auto-fluorescent based on deviations from the reference initial fluorescence values, as described in the Methods section **(Figure 2A)**. Compounds whose resulting Fnorm ‰ values exceeded three standard deviations above the reference mean were designated as potential hits. This yielded 54 candidates from the 653-compound library. After excluding quenchers and auto-fluorescent compounds, the final hit set was reduced to 37 molecules, corresponding to a 5.7% hit rate **(Figure 2B)**.

**Figure 2.**
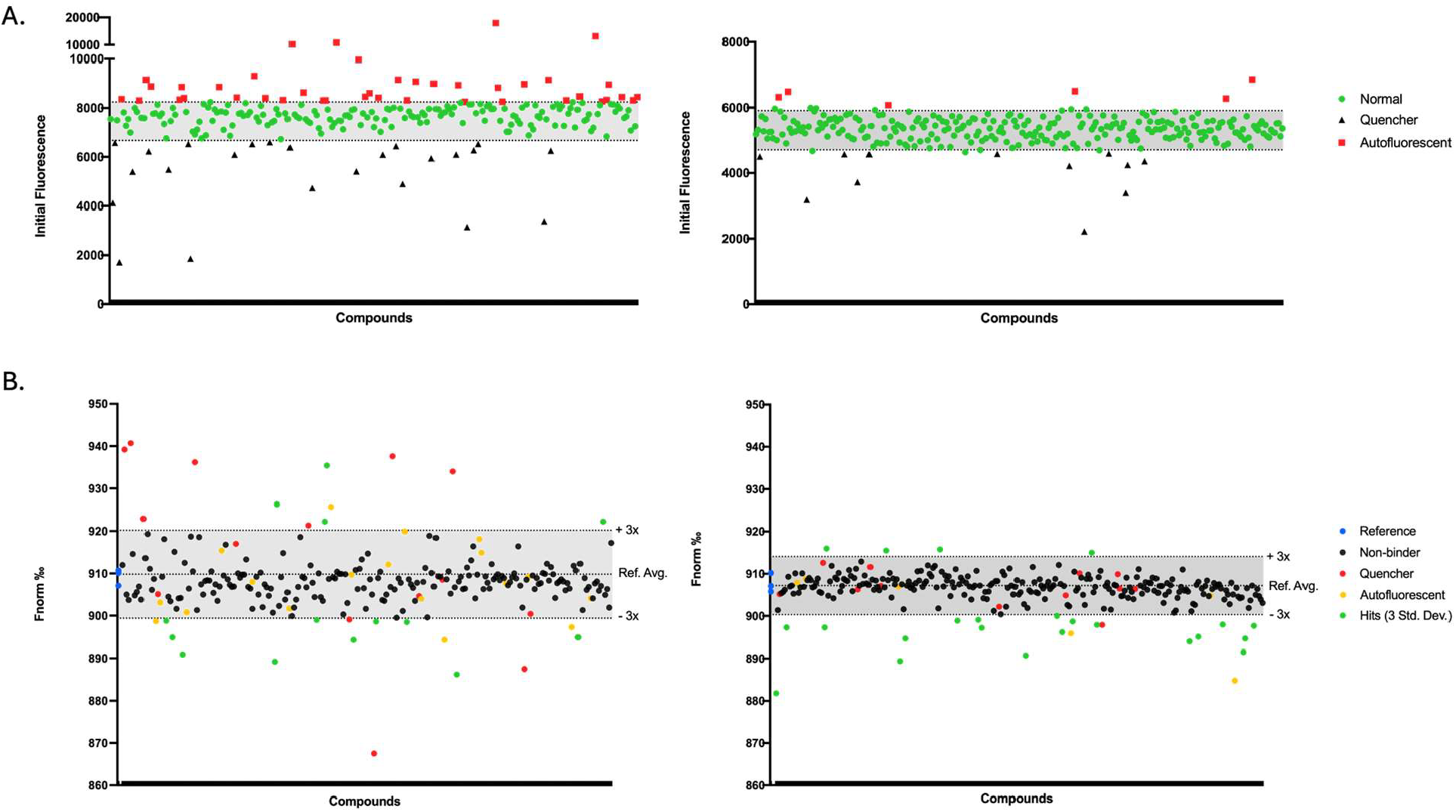
TargetMol Lipid Metabolism library screening results. (**A**) Initial fluorescence readout showing compounds found to modulate fluorescence measurements. (**B**) Fnorm ‰ values of SLIT2 protein incubated with compounds, plate 1 (left) and 2 (right). Multiples of standard deviation and average reference Fnorm ‰ value are denoted with horizontal lines.

Out of the 37 compounds tested **(Table 1)**, bexarotene **(Figure 3A)** stood out by exhibiting a clear, dose-dependent binding interaction with SLIT2, yielding a dissociation constant (*K*_D_) of 2.62 µM **(Figure 3B)**.

**Table 1.**
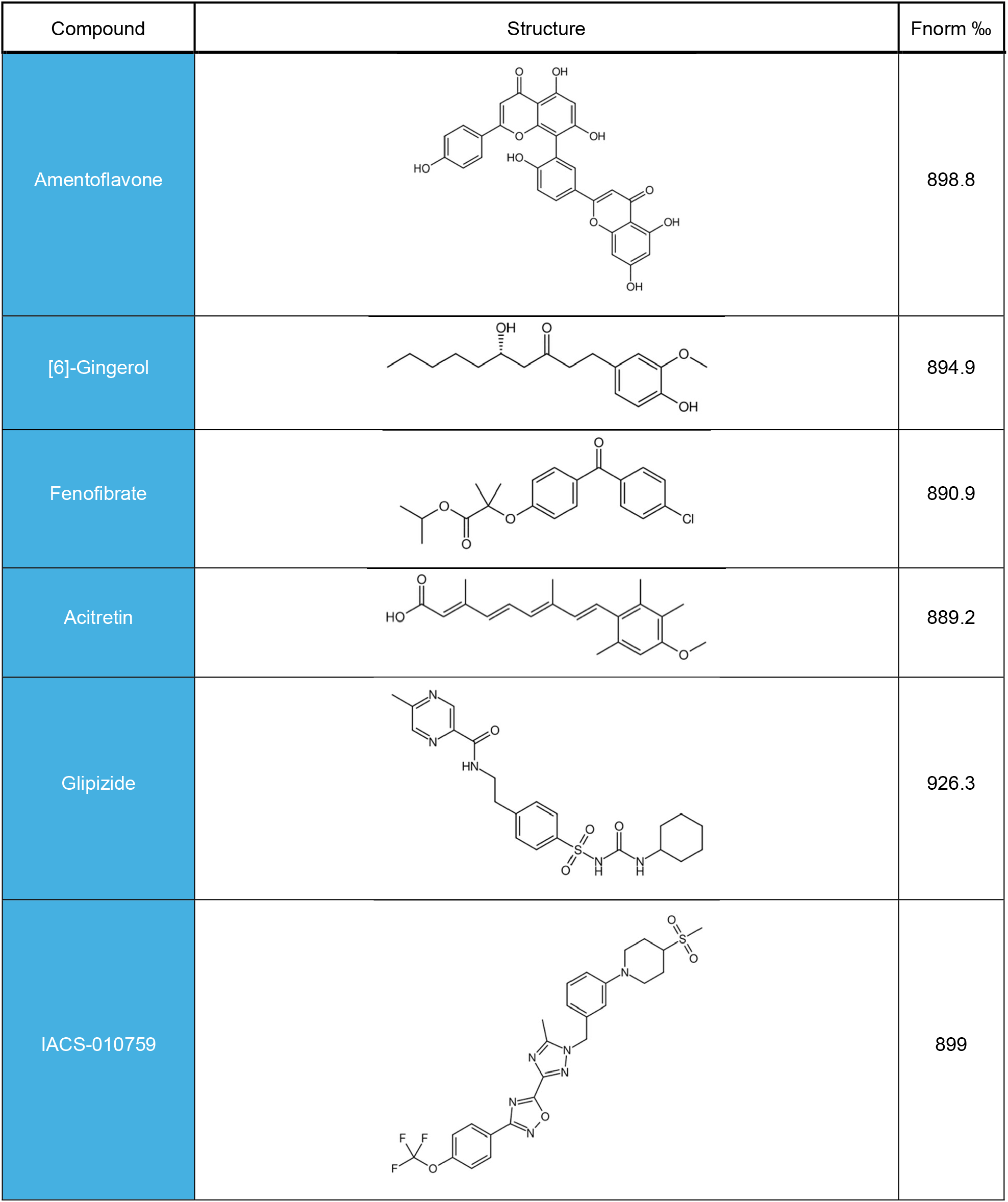
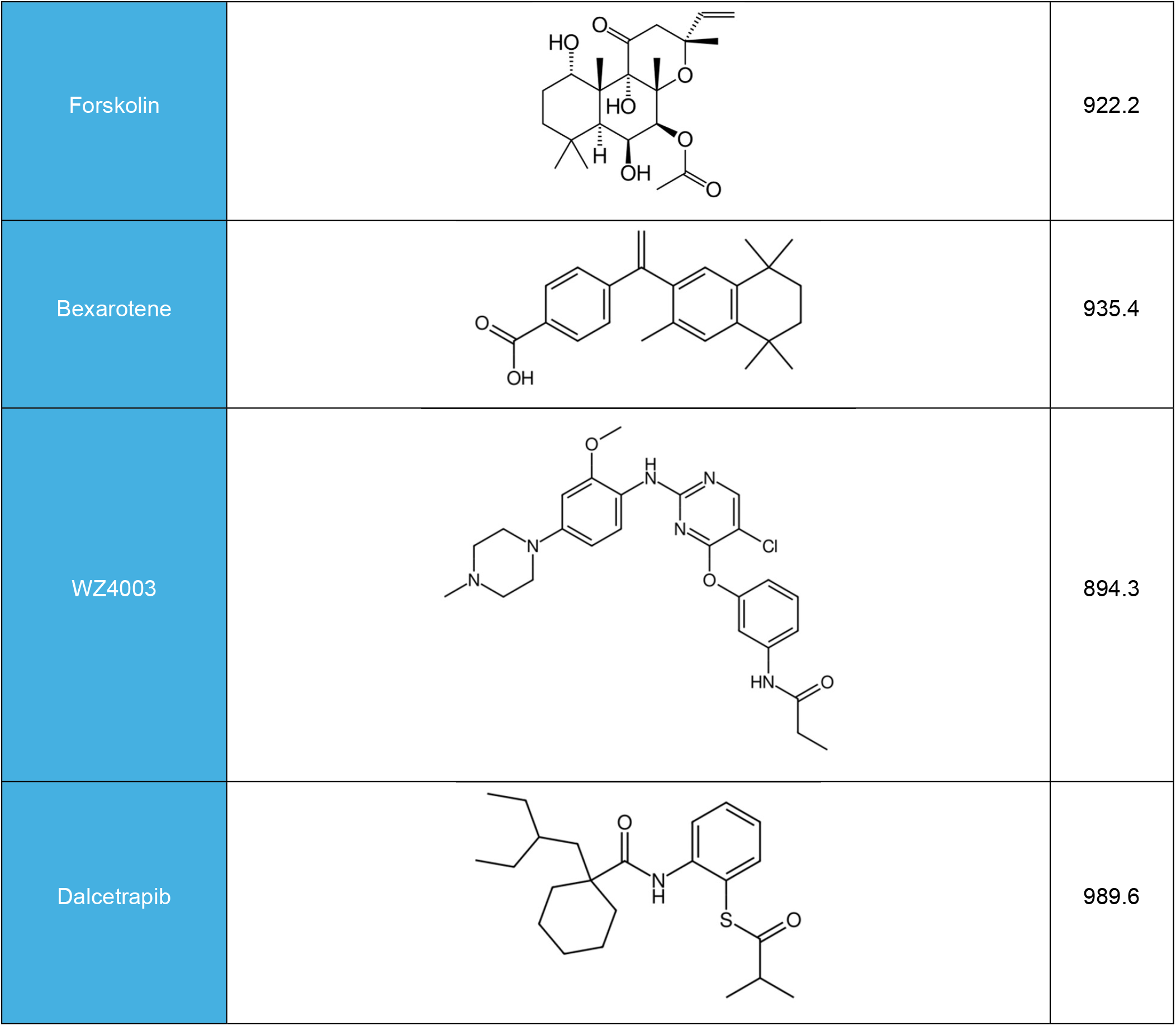
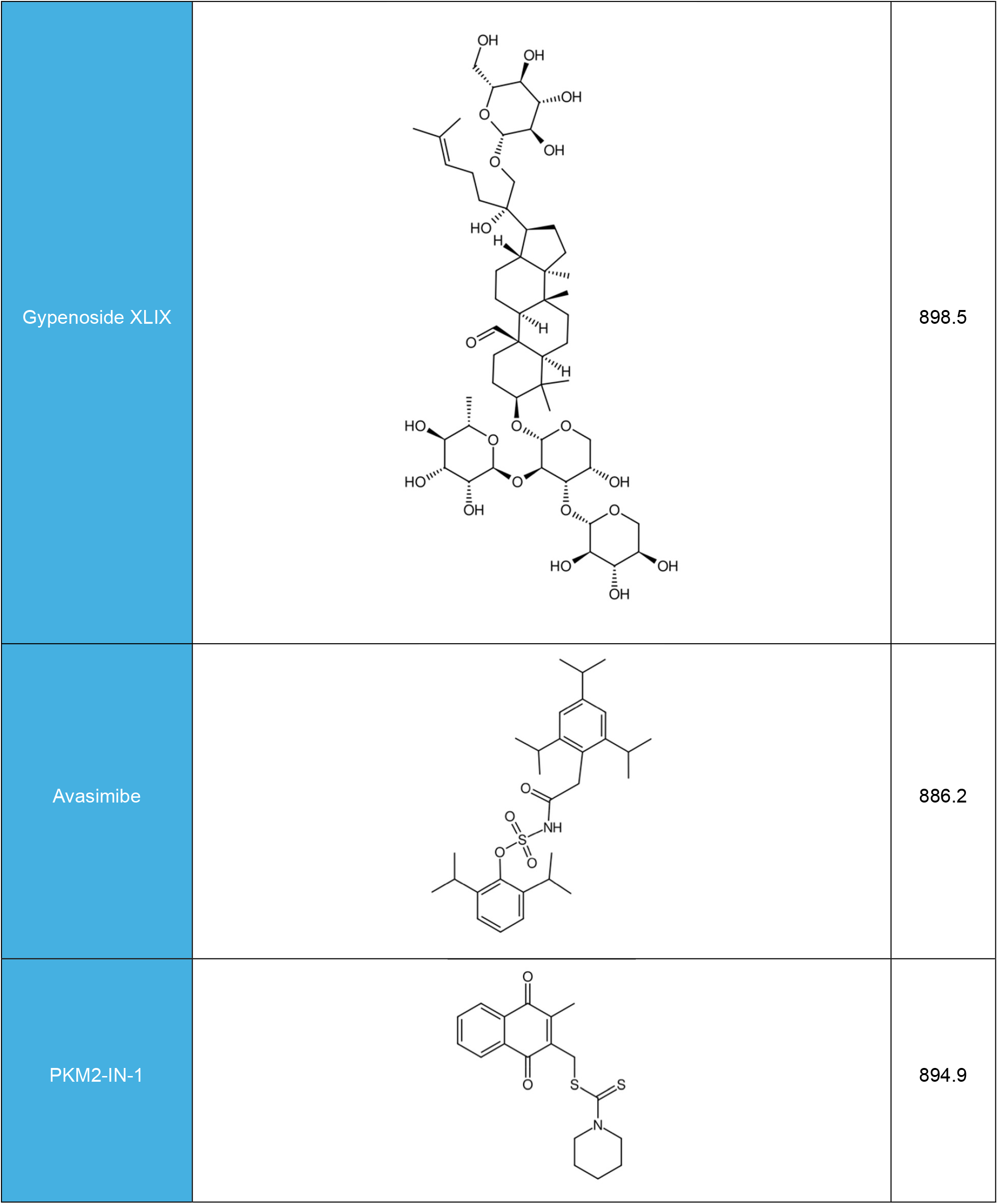
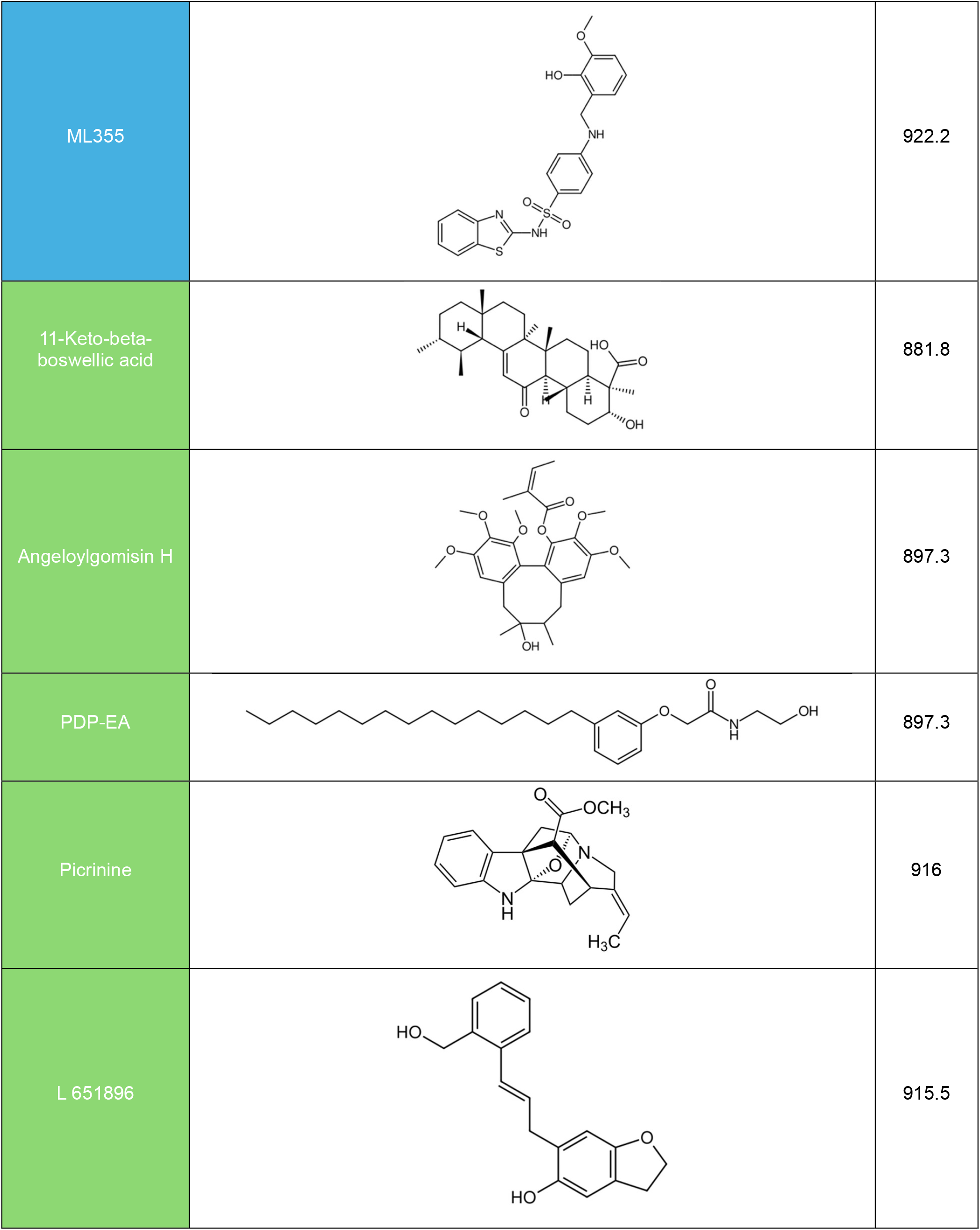
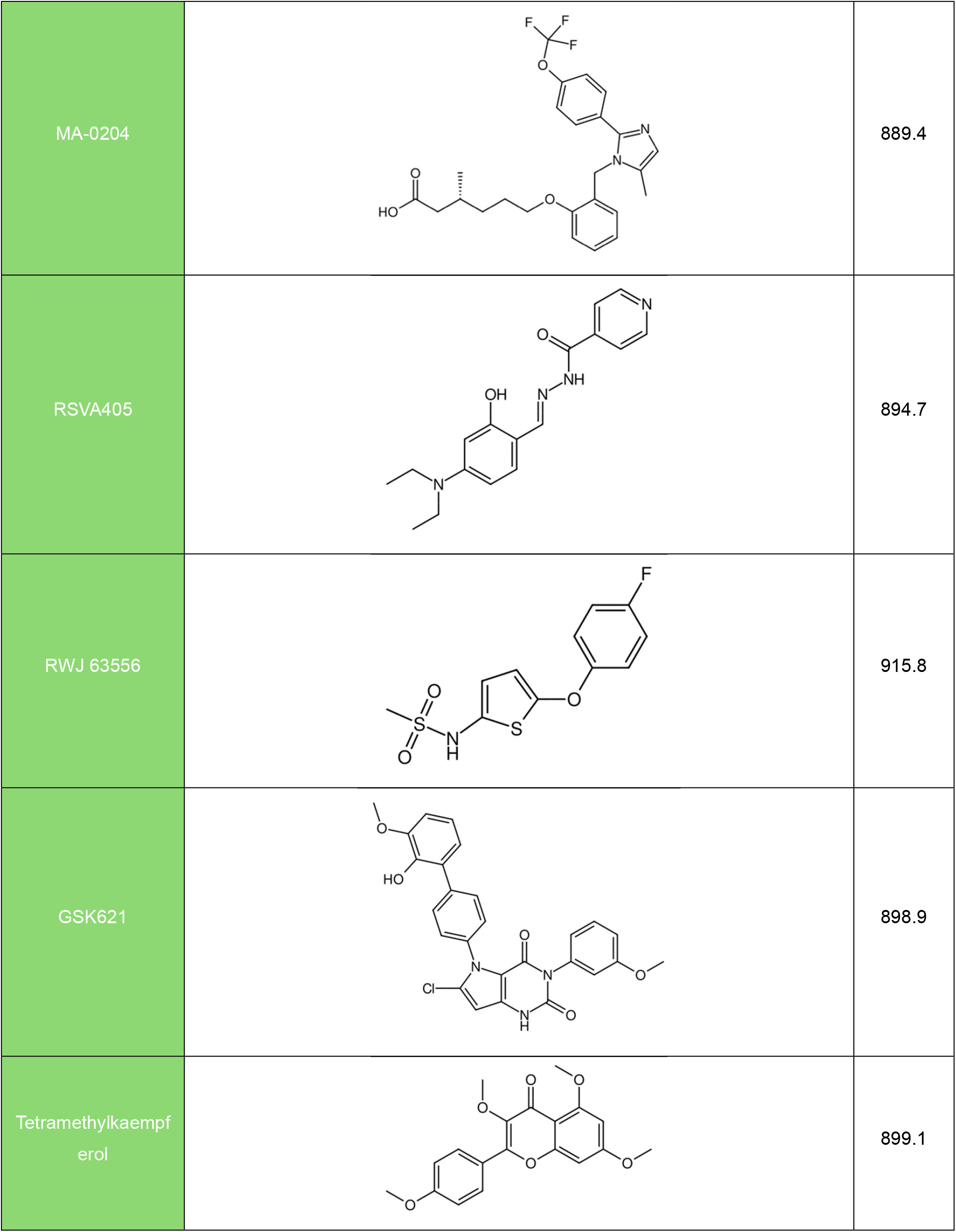
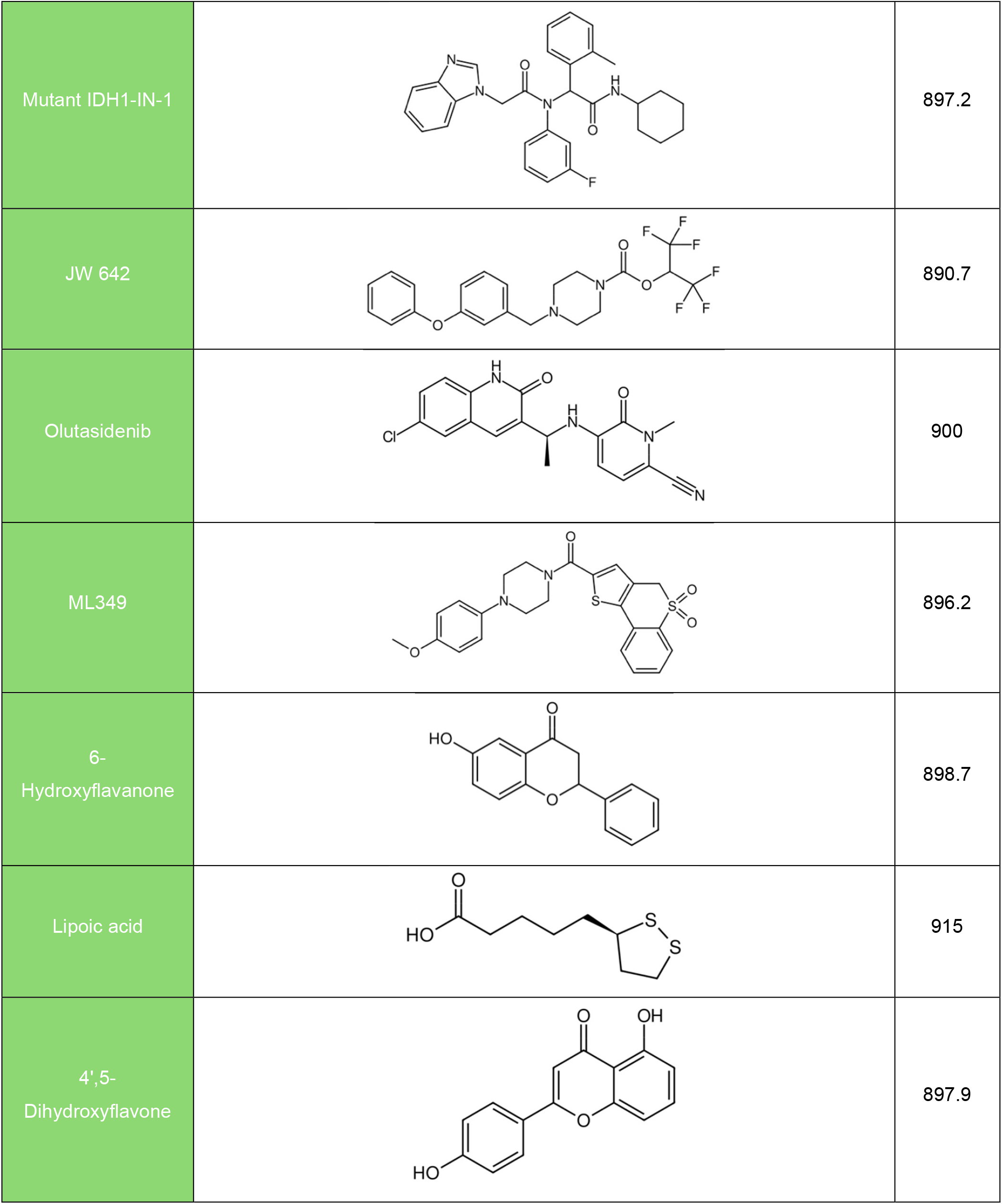
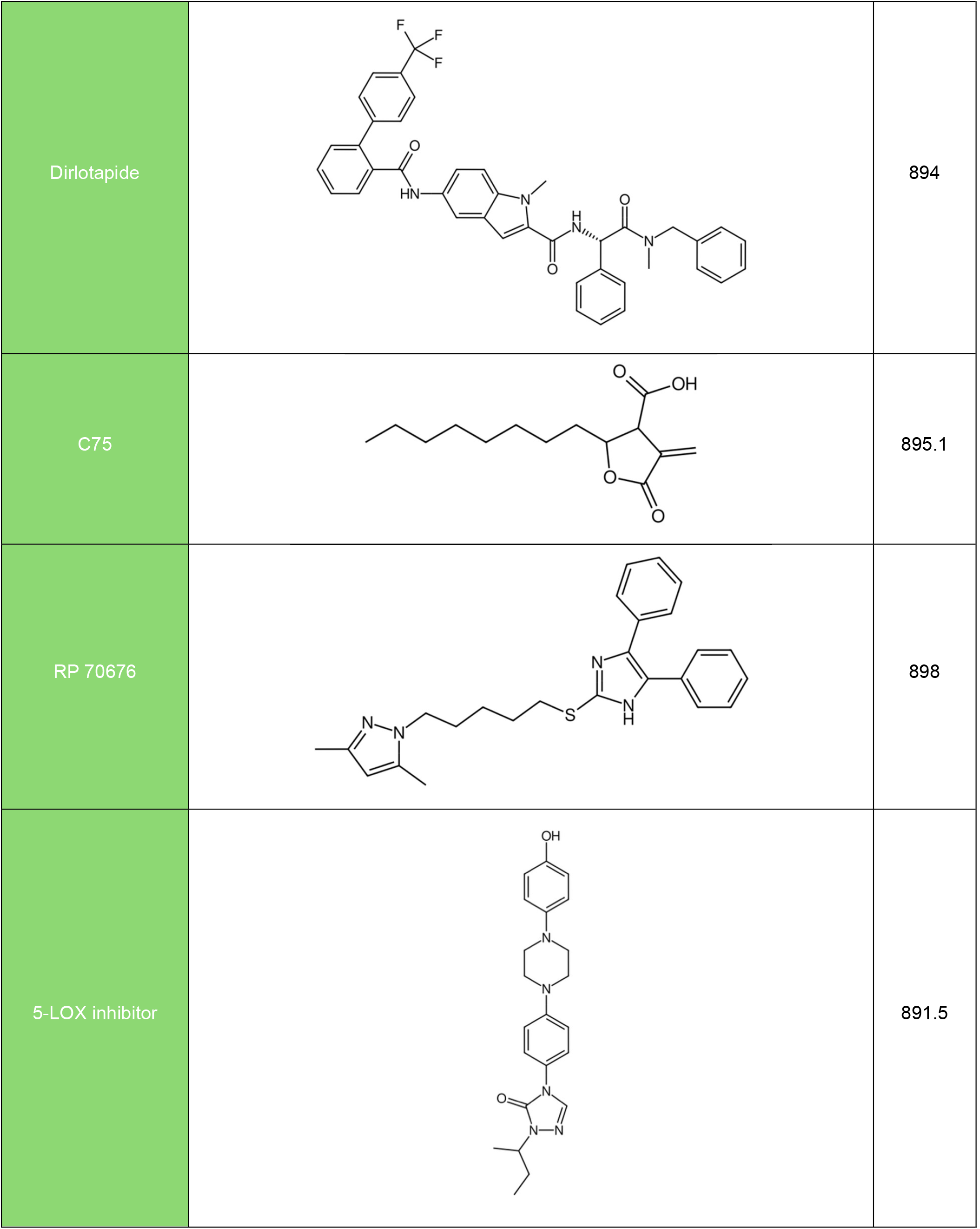
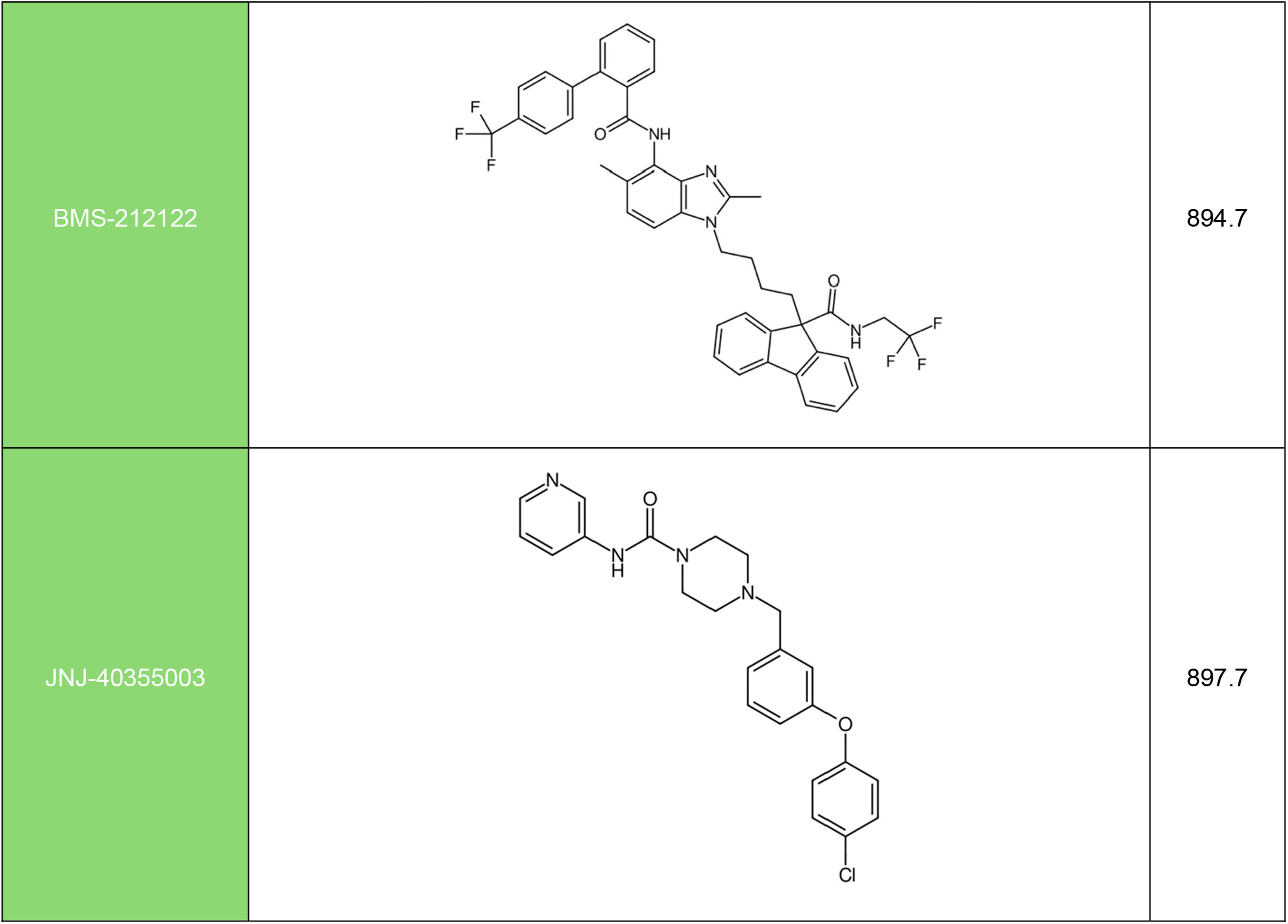
Structures, names, and Fnorm ‰ values of resulting 37 preliminary hits in Lipid Metabolism Library. Compounds in Plate 1 are labeled in blue, while those in Plate 2 are labeled in green.

**Figure 3.**
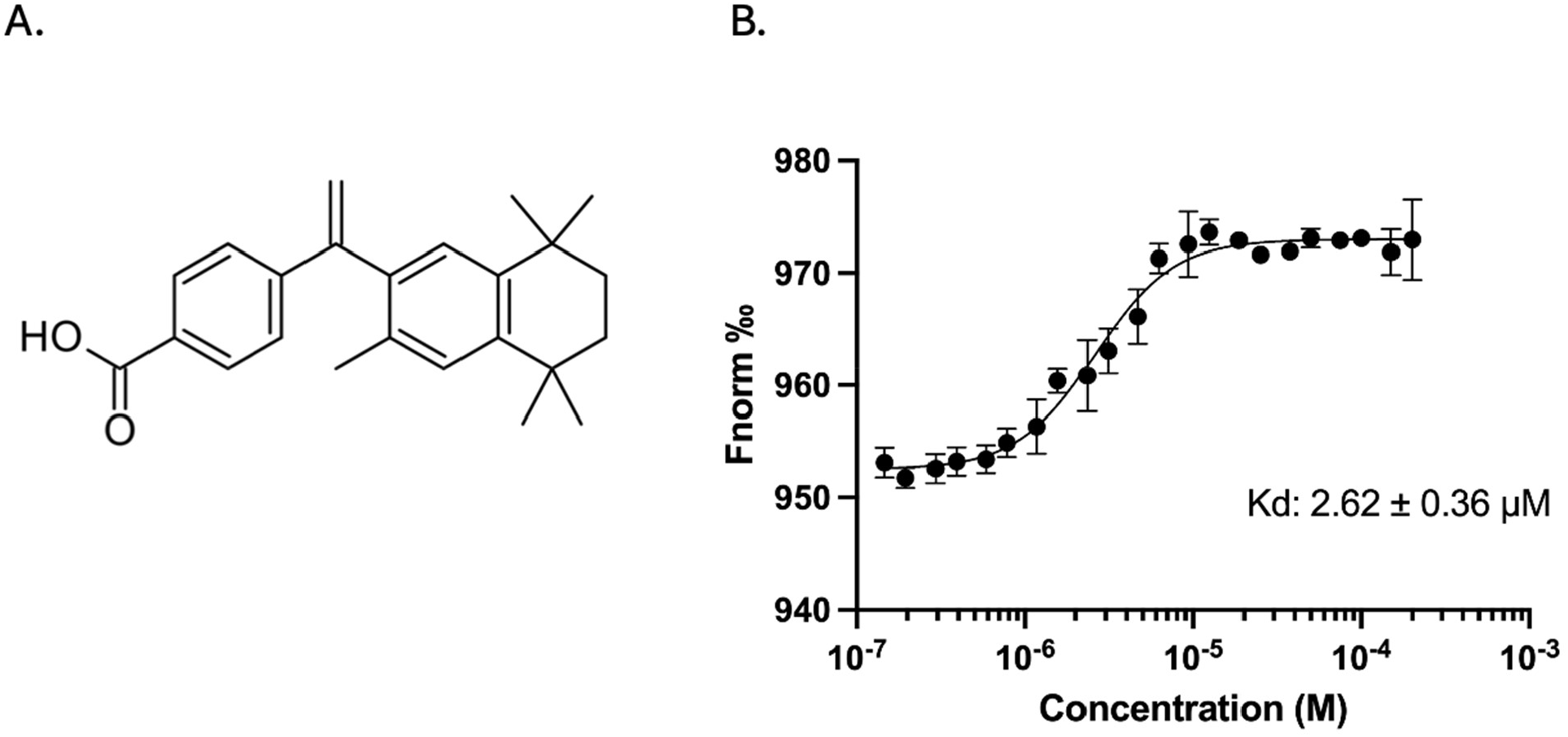
**(A)** Chemical structure of bexarotene. **(B)** Dose response binding curve of bexarotene for SLIT2 binding. Binding affinity results are shown as Fnorm ‰ vs Concentration (M) with the X-axis displayed on a log_10_ scale.

To validate our screening approach, we tested bexarotene in a TR-FRET assay designed to assess direct modulation of the SLIT2/ROBO1 interaction (19). Bexarotene demonstrated a consistent, dose-dependent reduction in TR-FRET signal, with an apparent IC_50_ of ∼22.8 µM and maximal inhibition around 15-25% (**Figure 4**).

**Figure 4.**
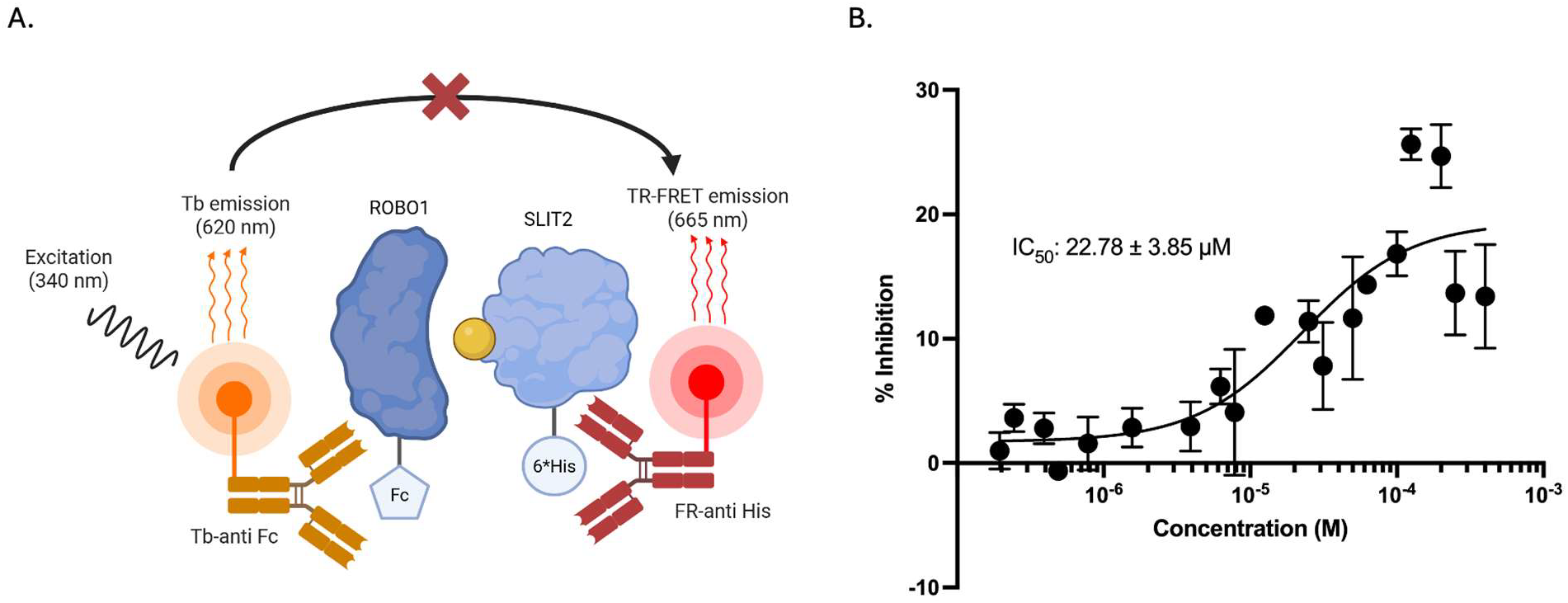
SLIT2 inhibition activity of Bexarotene. **(A)** Scheme of inhibition of SLIT2/ROBO1 TR-FRET signal by a SLIT2 inhibitor. **(B)** SLIT2/ROBO1 FRET inhibition dose response of Bexarotene. Concentration range of 400µM-1.95nM results in a maximal inhibition of ∼15-25% and an IC_50_ of 22.78 ± 3.85 µM.

## 4. Discussion

In this study, we present a scalable screening strategy for the identification of small molecule SLIT2 binders, integrating TRIC technology with TR-FRET-based validation. Our proof-of-concept screening resulted in the identification of bexarotene as a small molecule SLIT2 binder, with a *K*_D_ of 2.62 µM, with a maximal inhibition of ∼15-25% and an IC_50_ of 22.8 µM.

Bexarotene is a synthetic rexinoid that binds with high affinity to retinoid X receptors (RXRα, β, and γ), key nuclear receptors that regulate gene expression programs involved in differentiation, apoptosis, and lipid metabolism. Originally developed as an anti-cancer agent, bexarotene has a *K*_D_ in the low nanomolar range—approximately 14 nM for RXRα, 21 nM for RXRβ, and 29 nM for RXRγ—indicating strong and selective engagement across RXR subtypes (16,17). This tight binding allows it to effectively drive RXR-dependent transcriptional activity, which is central to its approved use in treating cutaneous T-cell lymphoma. Beyond oncology, its pharmacological profile has spurred interest in diseases linked to lipid dysregulation and neurodegeneration, although clinical efficacy outside of cancer remains limited. Importantly, its on-target effects often manifest systemically, leading to predictable adverse events like hyperlipidemia and hypothyroidism due to downstream RXR activity in metabolic and endocrine tissues (18).

Bexarotene binding to SLIT2 in the micromolar range is intriguing given its primary role as a nuclear RXR agonist. Its lipophilic nature may allow it to interact with hydrophobic surfaces, and SLIT2’s large extracellular domains likely provide such binding sites. Notably, the ability of bexarotene to disrupt the SLIT2/ROBO1 interaction in the TR-FRET assay supports the validity of the screening platform in identifying functional modulators. Importantly, these results illustrate that the combination of TRIC screening with FRET- based validation constitutes a viable platform for targeting the SLIT2/ROBO1 axis. This scalable assay will facilitate future larger screening studies and may ultimately contribute to therapeutic strategies. Beyond SLIT2, this dual biophysical screening approach can be readily adapted to other extracellular or “undruggable” protein–protein interactions, including guidance molecules such as CHI3L1, semaphorins, and netrins. The combination of TRIC and TR-FRET offers a versatile, label-compatible strategy for probing large, multidomain glycoproteins in a high-throughput format. As such, the platform described here provides a generalizable framework for extracellular target discovery where traditional screening approaches remain limited.

While this study establishes a robust biophysical screening platform for SLIT2-targeted small molecules, several limitations should be acknowledged. First, the functional consequences of SLIT2 binding by bexarotene were not evaluated in cellular or in vivo models. Follow-up studies are needed to determine whether such binding translates into modulation of SLIT2-driven signaling or immune phenotypes in relevant biological systems. Second, the screening was limited to a relatively small, lipid-focused compound library, and additional screening with chemically diverse or larger libraries may yield more potent or selective modulators. Third, although the TRIC and TR-FRET assays provide orthogonal validation of binding and functional disruption, further structural or mechanistic studies—such as docking, mutagenesis, or NMR—are warranted to define the interaction site and binding mode of bexarotene. Despite these limitations, the current platform lays important groundwork for systematic SLIT2-targeted discovery and can be readily adapted for other extracellular protein–protein interactions previously considered undruggable.

## Acknowledgments

We acknowledge funding support by R01CA293456 (PI Gabr) from the National Cancer Institute (NCI).

